# Identification of *cis*-regulatory mutations generating *de novo* edges in personalized cancer gene regulatory networks

**DOI:** 10.1101/130641

**Authors:** Zeynep Kalender Atak, Hana Imrichova, Dmitry Svetlichnyy, Gert Hulselmans, Valerie Christiaens, Joke Reumers, Hugo Ceulemans, Stein Aerts

**Affiliations:** Laboratory of Computational Biology, Center for Human Genetics, University of Leuven, Leuven, Belgium; Janssen Pharmaceutica NV, a Belgian company with registered offices at Turnhoutseweg 30, 2340 Beerse

## Abstract

The identification of functional non-coding mutations is a key challenge in the field of genomics, where whole-genome re-sequencing can swiftly generate a set of all genomic variants in a sample, such as a tumor biopsy. The size of the human regulatory landscape places a challenge on finding recurrent *cis*-regulatory mutations across samples of the same cancer type. Therefore, powerful computational approaches are required to sift through the tens of thousands of non-coding variants, to identify potentially functional variants that have an impact on the gene expression profile of the sample. Here we introduce an integrative analysis pipeline, called *μ- cis*Target, to filter, annotate and prioritize non-coding variants based on their putative effect on the underlying 'personal' gene regulatory network. We first validate *μ-cis*Target by re-analyzing three cases of oncogenic non-coding mutations, namely the *TAL1* and *LMO1* enhancer mutations in T-ALL, and the *TERT* promoter mutation in melanoma. Next, we re-sequenced the full genome of ten cancer cell lines of six different cancer types, and used matched transcriptome data and motif discovery to infer master regulators for each sample. We identified candidate functional non-coding mutations that generate *de novo* binding sites for these master regulators, and that result in the up-regulation of nearby oncogenic drivers. We finally validated the predictions using tertiary data including matched epigenome data. Our approach is generally applicable to re-sequenced cancer genomes, or other genomes, when a disease- or sample-specific gene signature is available for network inference. *μ-cis*Target is available from http://mucistarget.aertslab.org.

## Introduction

Oncogenic programs are characterized by aberrant gene expression profiles. A gene regulatory network underlying a cancer transcriptome can be considered as a perturbed stable network configuration, or as a cancer attractor state [1]. Gene expression changes leading from a normal cell to a malignant state are generally due to a series of acquired somatic mutations, which often affect proteins playing a key role in transcriptional regulation [2]. These can include mutations, amplifications, or translocations leading to an altered function or expression of transcription factors (e.g., MYC, TAL1, MITF, TP53), co-factors (EZH2, RB1, IDH1, MLL), or signalling molecules that lead to downstream alterations in transcription factor activity (e.g., RAS/RAC/RAF, KIT, PTEN, CDKN2A). More subtle changes can also occur in gene regulatory networks, which may cause fine-tuning of the emerging transcriptome rather than necessarily yielding a different attractor state. Such local network changes can involve the addition or removal of an edge in the network, affecting a single interaction between a transcription factor and a target gene. Edge perturbations can be caused by a mutation of a transcription factor binding site in a promoter or enhancer, leading to a *de novo* gain, or a loss, of the binding site, and a consequential expression change of a nearby target gene. Several examples of such perturbations are known to be associated to oncogenic programs, such as the gain of an ETS-family binding site in the *TERT* promoter, the gain of a MYB binding site in a 7.5 kb upstream *TAL1* enhancer [3–5], and the recently identified gain of a MYB binding site 4 kb upstream of *LMO1* oncogene [6]. Note that whereas these three examples occur recurrently across melanoma or liver cancer (for *TERT*) or across T-cell acute lymphoblastic leukemia (for *TAL1* and *LMO1*), they represent exceptional cases, since whole-genome sequencing, even across large cohorts such as 560 breast cancer genomes [7] failed to identify additional binding site changes that are significantly recurrent [8] (see also recent reviews [9–11]). This suggests either that *cis*-acting mutations are usually passenger mutations rather than driver mutations, or that they can occur as drivers at diverse positions, spread across hundreds of kilobases affecting the regulation of a target gene. The latter would render current cohort sizes underpowered, and would require different approaches to identify causal *cis*-regulatory mutations and their downstream consequences.

Computational predictions of a gain or loss of a transcription factor binding site can be performed by scoring the reference and mutated sequence with a position weight matrix (PWM) of the candidate factor [12,13]. This results in a “delta” PWM score, to which an arbitrary threshold can be applied to decide whether the gain or loss is strong enough. Such an approach is implemented in various bioinformatics tools, such as FunSeq2 [14] and OncoCis [15]. However, as position weight matrices are notorious in producing false positive predictions, also the delta score results in an excess of false positive gains or losses of binding sites. A possible solution to this problem is to take the context of the binding site into account, thus the encompassing regulatory region (promoter or enhancer). For example, the gain of MYB binding site in a random genomic position may not lead to *de novo* enhancer activity, whereas such a gain in the context of RUNX binding sites (MYB and RUNX bind together to leukemia enhancers) may cause ectopic enhancer activity [16]. Computationally, this solution depends on training more complex enhancer models, for example based on k-mer SVMs [17], Random Forests [16], or deep learning (deepSEA [18]). The main limitation of this approach is the dependence on high-quality training data to construct accurate enhancer models [16].

Here we investigate how “personalised” gene regulatory network reconstruction can be used to identify specific candidate *cis*-regulatory driver mutations in cancer genomes. Gene regulatory network inference is a common technique that has provided insight into master regulators in many cancer types [19–21], and the targets they regulate. Here, we exploit gene regulatory networks for the prioritization of non-coding mutations. Particularly, by first identifying the master regulators operating in cancer sample, we can identify those non-coding mutations that generate *de novo* targets of these master regulators. We develop an online tool to streamline this process, called *μ-cis*Target, and we demonstrate the use of *μ-cis*Target on known cases of *TERT* promoter and *TAL1* enhancer mutations. Finally, we predict new *cis*-regulatory mutations in ten cancer cell lines for which we sequenced the genome, transcriptome, and epigenome.

## Results

### A small number of non-coding mutations generate *de novo* oncogenic edges in driver gene regulatory networks

We developed a new computational pipeline, called *μ-cis*Target, to identify *cis*-regulatory mutations in a cancer sample, when both the whole genome sequence and the gene expression profile of that sample are available. The concept behind *μ-cis*Target is to simultaneously identify “personalized” candidate master regulators for a given cancer sample, based on the gene expression profile of the sample (and optionally combined with a 'general' cancer gene signature of the same cancer subtype), and to prioritise SNVs and INDELs in the non-coding genome of the sample by their likelihood to generate *de novo* binding sites for any of these master regulators (**Figure 1**). Among the list of candidates, we further determine a final set of mutations by applying two filters, namely: (i) the transcription factor for which a binding site is generated is itself expressed in the sample and is related to the cancer type; and (ii) the mutation is located close (up to 1 megabase (Mb)) to a target gene that is over-expressed, and within the same topologically associating domain (TAD) as the over-expressed target, it is related to the cancer type under study, and/or it is a potential driver gene (**Figure 1**, see Methods). These criteria are largely inspired by previously published *cis*-regulatory driver mutations, such as those driving *TERT*, *TAL1* and *LMO1* [3–6]: these oncogenes are over-expressed, and the generated binding sites are bound by (over-) expressed and cancer-type relevant transcription factors, namely GABPA for *TERT* and MYB for both *TAL1* and *LMO1*.

**Figure 1.**
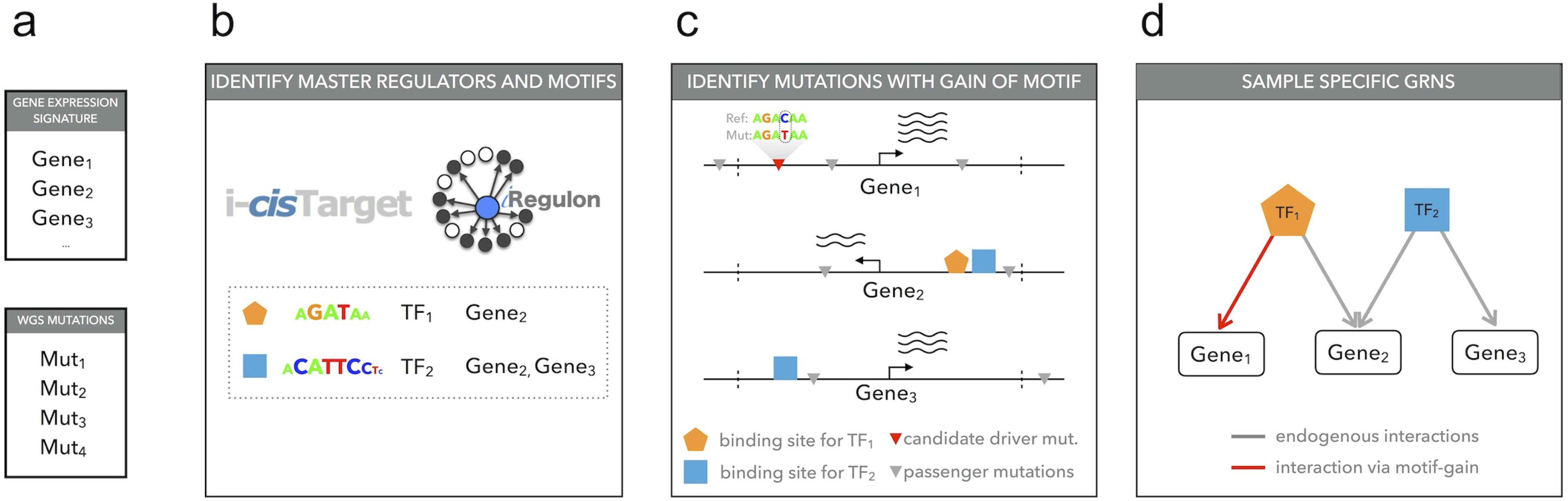
Overview of the *μ-cis*Target pipeline to predict *cis*-regulatory mutations. **a)** As input *μ-cis*Target takes a gene signature and a list of genomic variations. The gene signature can be derived from the matched transcriptome of the same cancer sample, or can be a general gene signature of the matching cancer subtype. **b)** Motif discovery on the gene signature yields enriched motifs and candidate transcription factors. Motif discovery can be performed using i-*cis*Target or iRegulon. **c)** Variations are selected by their proximity (<1Mb) from the genes in the input gene signature, and are scored with the motifs found under b). **d)** Genes with gains of motifs for cancer-type-related factors that are expressed in the sample are added to the inferred gene regulatory network (red edge). An optional filtering step selects only over-expressed cancer-related driver genes as targets.

To illustrate how *μ-cis*Target works we first apply it to a simulated set of 67 variants spread around the *TAL1* gene (up to 1 Mb upstream or downstream), where we inserted the true driver variant that generates a *de novo* MYB binding site. In the first step of the method, we used as input the top 500 MYB ChIP-seq peaks obtained in the same sample where the variant occurs (the JURKAT cell line), which finds the MYB and RUNX1 motifs as enriched (**Supplementary Figure 1**). This analysis thus infers a candidate network with MYB and RUNX1 as master regulators (**Figure 2a**). Among the 68 variants, only one generates a new binding site for any of the enriched motifs, which is the true driver mutation, with a strong gain for the MYB motif (**Figure 2b-c**). We then used the same master regulators to interrogate the recently discovered *LMO1* enhancer mutation and again we could correctly predict MYB gain of motif as a result of this non-coding mutation (**Figure 2b,d**).

**Figure 2.**
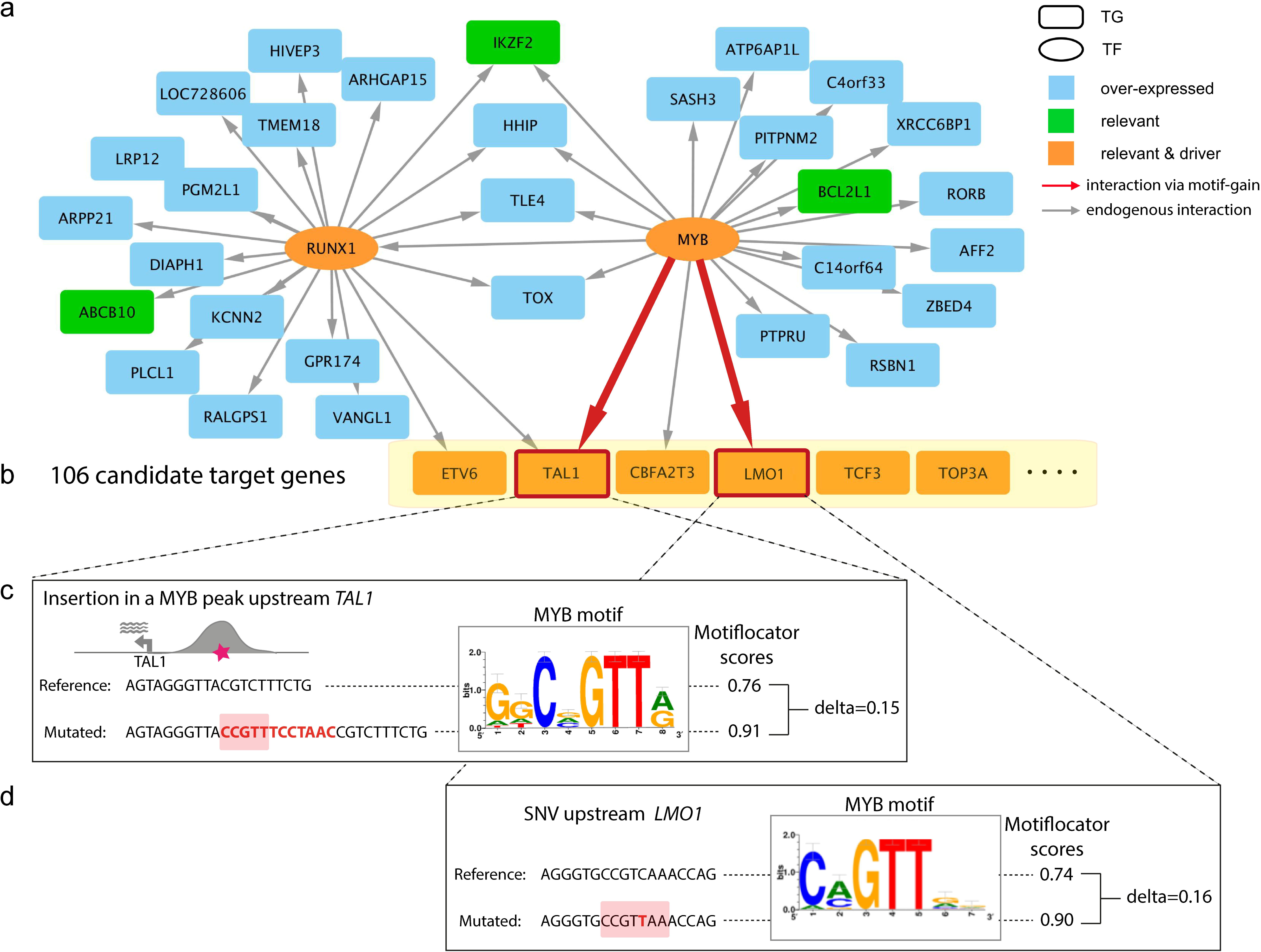
Detection of *TAL1* insertion and *LMO1* mutation in JURKAT cell line. **a)** Gene regulatory network inferred from the top 500 MYB ChIP-seq peaks on JURKAT cell line (by i-*cis*Target [62]). The top enriched motifs are directly annotated for RUNX1 and MYB transcription factors (TFs), which are also expressed in the JURKAT cell line. Only over-expressed target genes (TGs) in JURKAT are shown (blue nodes) from which some of them are moreover relevant to leukemia cancer type (green nodes) and some are known as cancer drivers (orange nodes). The grey edges represent the link between the TF and TG based on the presence of the TF motif in a MYB ChIP-seq peak near (<1Mb) the target gene. **b)** Non-coding mutations close to candidate target genes that are over-expressed, relevant and drivers are tested by MotifLocator to find candidate mutations that yield a motif gain. Because there is no whole genome available for JURKAT cell line, we simulated a dataset with the JURKAT insertion upstream of *TAL1* together with 67 control mutations from 10 sequenced cancer cell lines (see **Table 1**), that are found in the *TAL1* locus. Out of all 68 mutations, only the JURKAT insertion showed the gain of MYB motif, which caused a new link between *TAL1* and MYB (red arrow). **c, d)** Details of the JURKAT insertion 7.5 kb upstream of the *TAL1* oncogene (c) and the JURKAT SNV 4 kb upstream of the *LMO1* oncogene (d), where the reference and mutated sequences are shown (the insertion/SNV in red, the core of the motif is highlighted) together with their scores given by MotifLocator for the master MYB motifs.

**Table 1.**
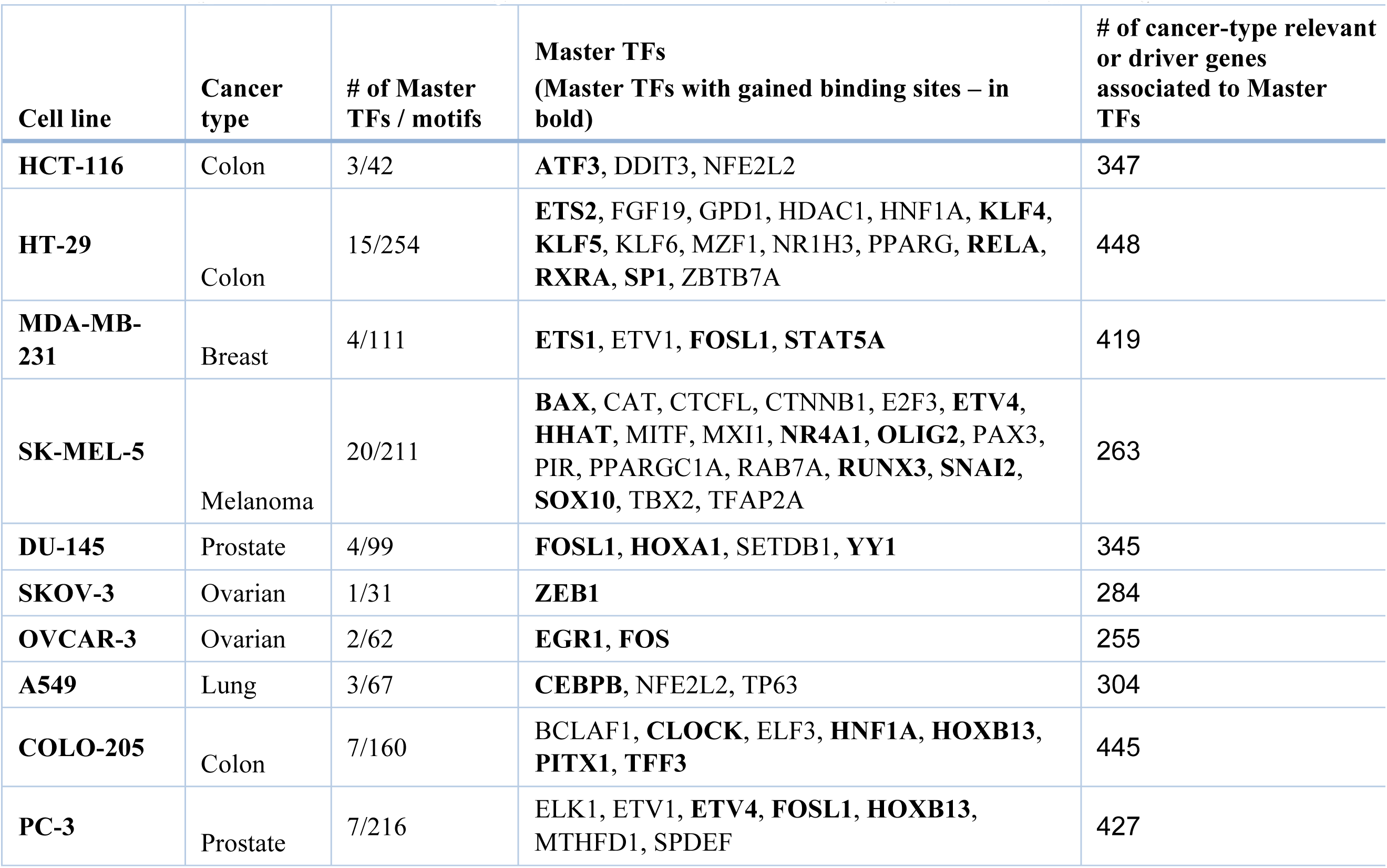
Sample-specific master regulators predicted from sample-specific gene signatures. The table lists the number of predicted master regulators per cell line (together with the number of motifs directly associated with these TFs; the names of the master TFs (bold indicating gains), and finally the number of candidate over-expressed cancer-type related or driver genes (near these genes we score mutations for motif gains).

Next, we tested whether *μ-cis*Target could identify the well-known *TERT* promoter mutation from a sample with a fully sequenced genome (having the *TERT* mutation) and a matched transcriptome. The *TERT* promoter mutation results in a *de novo* ETS-binding site and occurs in 55% of melanoma samples [22]. We selected 7 melanoma samples from TCGA with a *TERT* promoter mutation and for which expression and whole genome mutation data is available through TCGA. We ask whether *μ-cis*Target can identify the *TERT* promoter mutation in each individual sample as a candidate *cis* Gain-of-Function (*cis*-GoF) mutation, starting from the matched gene expression and mutation data. The first step consists of the identification of master regulators, starting from a gene signature of sample-specific up-regulated genes. For six out of the seven samples, *μ-cis*Target predicts at least one ETS family member as master regulator (**Supplementary Table 1**). The second step of *μ-cis*Target consists of identifying mutations that result in a gain-of-binding site near potential oncogenic drivers per sample (i.e. over-expressed genes that are either specific for the cell type or a potential driver gene, see **Supplementary Table 2**). In all those cases where an ETS factor is found as a master regulator, the *TERT* promoter mutations (both C228T and C229T) are predicted as gains of ETS binding sites (**Figure 3a-c**, **Supplementary Table 1**). Next, to test the specificity of our method we used whole-genome mutation calls for three of these melanoma samples and predicted candidate *cis*-GoF mutations (**Figure 3b**). From the initial 110K to 240K mutations, *μ-cis*Target identified 58 to 114 candidate *cis*-GoF mutations, including the *TERT* promoter mutations. All these candidates were either within introns or distal regulatory regions, while the *TERT* promoter mutations are among the few predictions located in a gene promoter (only TCGA-EE-A20H has two other candidate mutations that are located in a gene promoter) (**Supplementary Table 3**). This demonstrates that *μ-cis*Target is able to identify a manageable number of candidate functional non-coding mutations among thousands of candidates, while providing a prediction of their function in terms of sample-specific gene regulatory networks. More importantly, our results demonstrate that *μ-cis*Target can identify a functional non-coding mutation (such as the *TERT* promoter mutation) in a sample-centric manner without requiring recurrence across a large cohort.

**Figure 3.**
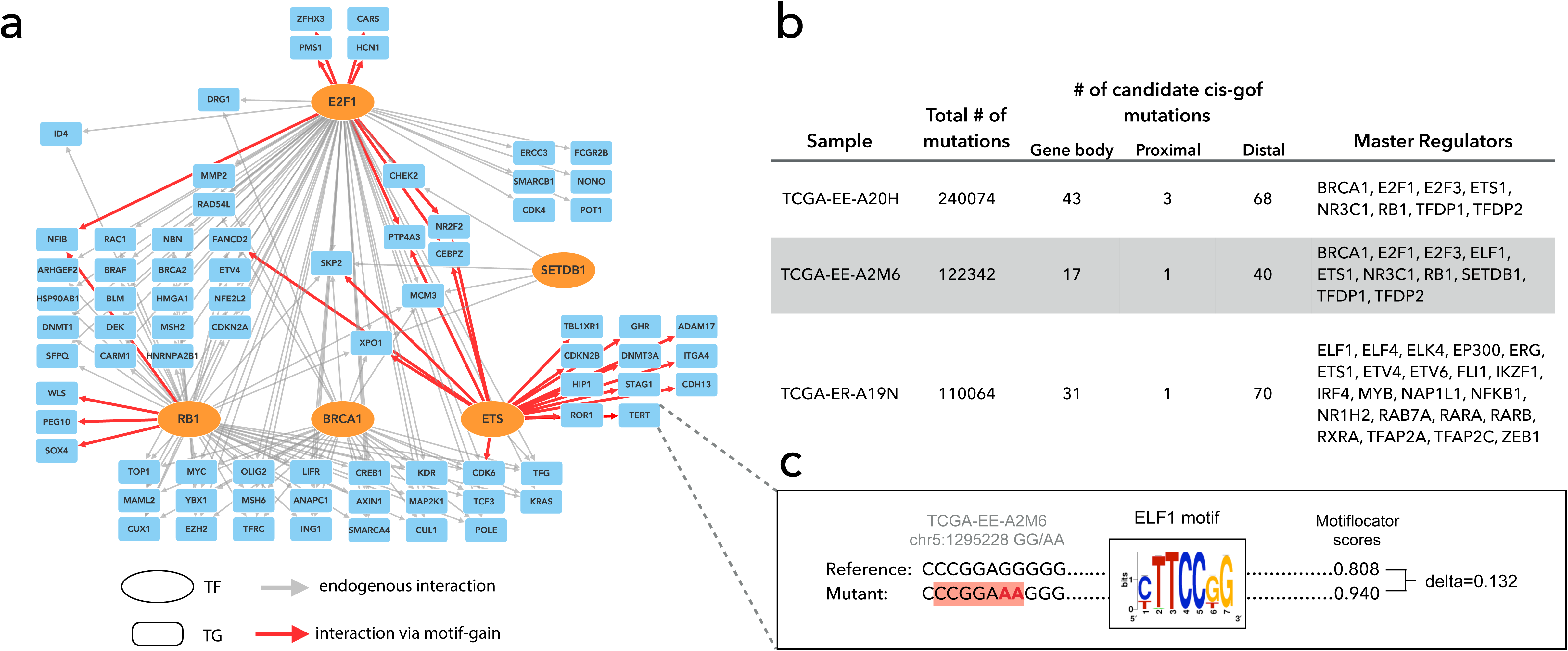
*TERT* mutation identification through personalized gene regulatory network of TCGA-EE-A2M6. **a)** Gene regulatory network inferred by iRegulon analysis from over-expressed genes (Z-score >=1) of melanoma sample TCGA-EE-A2M6. Among the enriched motifs are directly annotated motifs for ETS TFs (ELF1, ETS1), BRCA1, E2F family TFs (E2F1, E2F3, TFDP1, TFDP2), RB1, and SETDB1 (for simplicity the network is drawn only with cancer driver TFs, for a full list of predicted master regulators for this sample see Supplementary Table 1). The grey edges represent the link between TFs and TGs based on iRegulon analysis, while red edge indicates gain of ETS motif caused by the *TERT* promoter mutations C228T and C229T. All the represented TGs are over-expressed in TCGA-EE-A2M6, associated with melanoma and known cancer drivers. **b**) Detail of the mutation at the *TERT* promoter, where the reference and mutated sequences are shown (the mutation in red, the core of the motif is highlighted) together with their scores given by MotifLocator for the swissregulon__hs__ELF1_2_4.p2 motif (which is directly annotated to ELF1, ELF2 and ELF4).

### Application of *μ-cis*Target to ten re-sequenced cancer cell lines

After validating *μ-cis*Target on the *TAL1, LMO1,* and *TERT* mutations, we analysed ten widely used cancer cell lines as a discovery set (**Supplementary Table 4**). We essentially implemented the same strategy as in the validation cases. Namely, we first identify cell-line specific master regulators that are relevant genes per cell line using motif enrichment analysis. Next, we identify non-coding mutations in the genome of that cell line that create *de novo* binding sites for any of these master regulators, and that are near oncogenic drivers (relevant or driver genes (**Supplementary Table 2**)). For the first step we obtained gene expression profiles of these cell lines from COSMIC Cell Lines Project [23] and predicted master regulators using gene signatures of these cell lines (**Table 1**). For each sample, all genes expressed with Z-score above 1 (compared to all other cell lines in Cosmic Cell Line Project) are used for motif-enrichment based master regulator discovery. The initial set of predicted transcription factors are filtered for their own over-expression (Z-score above 1) and their cancer type specificity (based on **Supplementary Table 2**). Interestingly, our motif-based predictions of master regulators is supported by ChIP-seq data (when available) for 6 / 33 master regulators across the 10 cell lines (**Supplementary Table 4**).

Transcription factors identified at this step can be linked to several signalling pathways (**Supplementary Figure 2**) but two functional classes of transcription factors emerge at this step: lineage-associated transcription factors and EMT-associated transcription factors. Lineage associated transcription factors include MITF for the melanoma cell line SK-MEL-5 [24], TP63 for the lung cancer line A549 [25], KLF5 for the colon cancer line HT-29 [26], and ETS-family transcription factors for prostate, colon, ovarian and breast cancer cell lines [27–31]. This class of transcription factors is expressed at an earlier developmental stage and is reactivated during tumorigenesis. Another group of transcription factors are the EMT associated factors: FOSL1 for MDA-MB-231 and DU-145, ZEB1 for SKOV-3, FOS for OVCAR-3, and SNAI2 for SK-MEL-5. The majority of cell lines with these TFs as master regulators are derived from metastatic sites (SK-MEL-5, DU-145, PC-3). Of the remaining two cell lines, OVCAR-3 is derived from a chemo-resistant patient [32] and MDA-MB-231 demonstrates mesenchymal cell morphology [33] and is regarded as invasive *in vitro* [34]. Master regulators obtained at this step also corroborate well with what is known about these cell lines. For instance, the predicted master regulators for the lung cancer cell line A549 include NFE2L2 (NRF2) which is an essential gene for cell proliferation and chemoresistance in lung cancers, and specifically in A549 since knock-down of NFE2L2 in A549 inhibits proliferation [35]. Another example involves the MDA-MB-231 cell line for which ETS factors ETS1 and ETV1, as well as FOSL1 and STAT5A/B (the motif is directly annotated for both STAT5A and STAT5B) are found as master regulators. Gene knock-down studies involving these four transcription factors in this cell line demonstrated that each of these transcription factors are essential for growth, migration and metastatic potential of this cell line [36–39]. And lastly, it has been shown for the ovarian cancer cell line SKOV-3 that inhibition of ZEB1, which is predicted as a master regulator, hampers migration *in vivo* and tumor growth *in vitro* when xenografted in mice [40]. In conclusion, master regulator predictions seem to capture and represent oncogenic processes ongoing in these cell lines.

Next, we obtained whole genome mutation calls for 10 cell lines by re-sequencing them using a combination of Illumina and Complete Genomics (CG) technology (**Supplementary Table 5**). On average the cell lines contain 1.69 Mio variants (SNVs and INDELS combined) of which, on average, 98% are non-coding (**Table 1**, **Supplementary Table 5**). For each sample, we scored non-coding mutations using sample-specific master regulators identified in the first step. Again, we defined candidate *cis*-GoF mutations as variants that generate *de novo* binding sites for any of the predicted master regulators, near oncogenic drivers. There is a high variation between the number of candidate *cis*-GoF mutations between cell lines and this correlates with the number of *somatic* coding mutations for these cell lines (r = 0.96 & p-value < 0.05 except for HCT-116) (**Figure 4**, **Supplementary Figure 3**). Across all ten cell lines, *μ-cis*Target initially identifies 485 candidate mutations, and even though we assign mutations to genes in a regulatory space up to 1 Mb, almost all of them results in an association covered within a TAD (468/485). We focus on those 468 mutations associated to their targets within a known TAD affecting 290 oncogenic drivers (**Figure 4, Supplementary Table 6)**. Only 29 genes have a protein altering mutation (i.e. missense substitution, in-frame or frameshift indel) that might be associated with its over-expression and none of these genes are affected by copy number aberrations, thus 94% of the genes are affected only by non-coding mutations (**Supplementary Table 7**). Although there are no genomic positions recurrently mutated across the 10 cell lines, there are 24 genes that are recurrently affected by a *cis*-GoF mutation in 2 or more samples. For instance, FOXA1, which acts as pioneering factor in prostate cancer [41], is found be affected by *cis*-GoF mutations in the prostate cancer cell lines DU-145, PC-3 and in the colon cancer cell line HT-29 (**Supplementary Table 8**). Moreover, there are two *de novo* master regulator – target gene pairs recurrent across 10 cell lines: FOSL1- *FOXA1* in prostate cancer cell lines DU-145 and PC-3; and FOSL1- *MTRR* in the breast cancer cell line MDA-MB-231 and prostate cancer cell line DU-145. Although it may be possible that other binding site gains, located near genes that are not (yet) known as oncogenes for the cancer type under study, could play a role in the oncogenic program, we consider this as unlikely, given the large amounts of cancer type specific (and general) oncogenes that are known so far. In conclusion, *μ-cis*Target provides a short list of candidate *cis*-regulatory mutations, that have a potential impact on the expression of relevant oncogenes.

**Figure 4.**
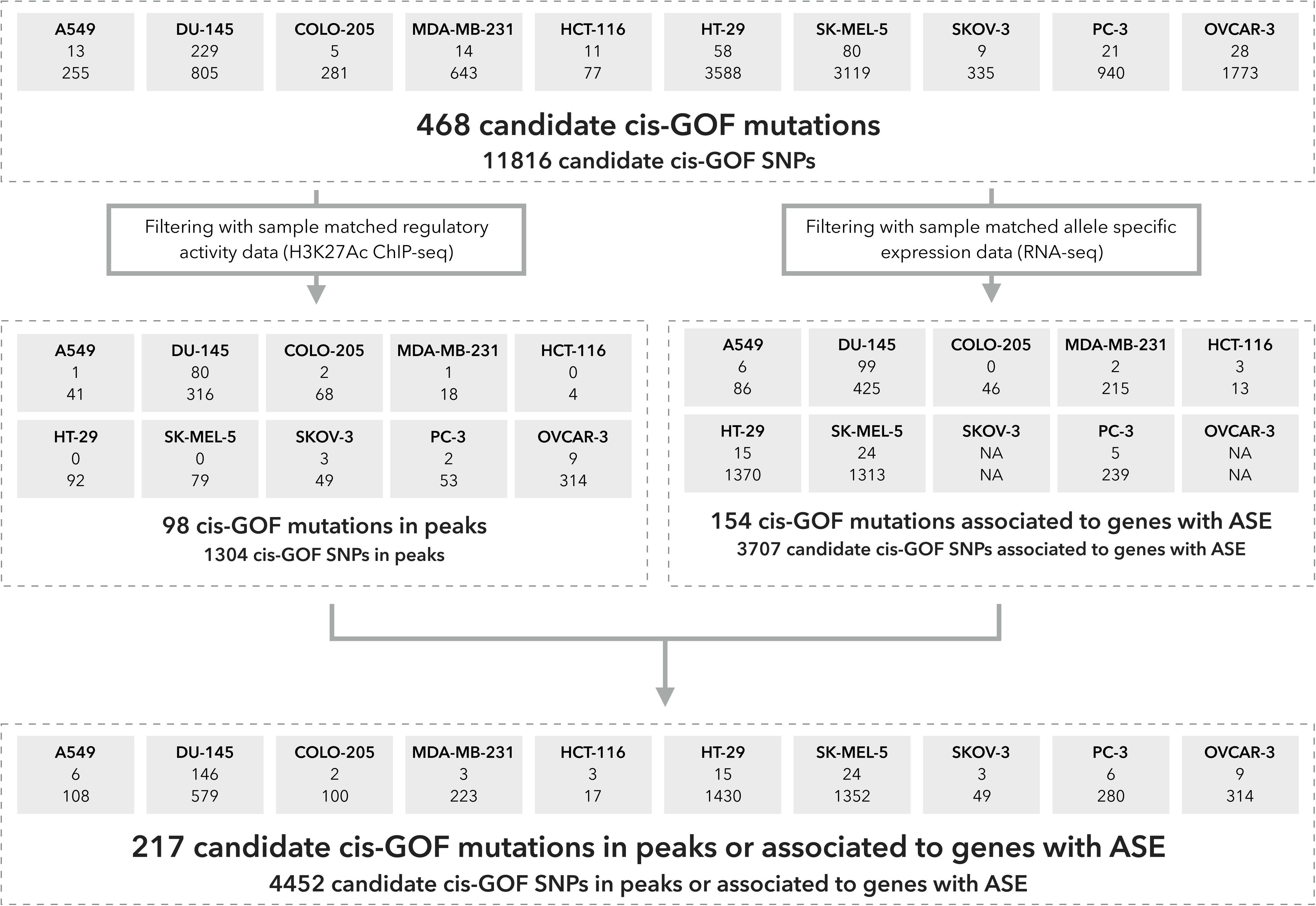
Candidate *cis*-GoF mutations in 10 cancer cell lines. For each cell line the number of candidate *cis*-GoF mutations (above) and SNPs (below) are indicated within each box. Across 10 cell lines we identified 468 candidate *cis*-GoF mutations and 11816 SNPs. Identified candidates are validated using matched regulatory (H3K27Ac ChIP-seq) and transcriptomic data, and overall 217 out of 468 candidate *cis*-GoF mutations (4452 out of 11816 SNPs) exhibited evidence for selection.

### Evaluation of predicted *cis*-regulatory mutations using matched epigenomes and allele-specific expression

Evaluating the potential impact of predicted oncogenic *cis*-mutations is challenging. Here, we test whether the predicted mutations may have an impact on the regulatory activity of the encompassing region. To this end, we use existing as well as newly obtained regulatory data, and allele specific expression information (as obtained from RNA-seq) for our 10 cell lines. Note that previous studies have used regulatory data to filter non-coding mutations [42], but these were not sample-matched. Here we explicitly use matched regulatory data for the same sample. Overlapping candidate *cis*-GoF mutations with sample matched H3K27Ac peaks revealed that 98 out of the 468 candidate mutations are in a potentially active regulatory region. For 6 out of the 10 cell lines, candidate *cis*-GoF mutations are enriched in active regulatory regions (hypergeometric test p-value <= 0.05, **Figure 4, Supplementary Table 9**). The same holds true for *cis*-GoF SNPs, since for all cell lines *cis*-GoF SNPs are enriched in active regulatory regions, indicating that *μ-cis*Target can identify potentially functional variants, be it SNPs or mutations. Additionally, we queried a large set of ChIP-seq peaks against other regulatory marks and transcriptions factors as obtained from the ChIP-Atlas database (http://chip-atlas.org) (433 datasets for 7 of our 10 cell lines, **Supplementary Table 10**, see Methods) which revealed an additional 12 *cis*-GoF mutations that are located in a TF ChIP-seq peak *in the corresponding sample* (**Supplementary Table 6**). In one of these examples, a predicted gain of a AP-1 binding site is observed upstream of the *RARB* gene in the breast cancer cell line MDA-MB-231, and this site co-localizes with a JUNB ChIP-seq peak (ChIP-seq performed in MDA-MB-231). Moreover, this mutation is indeed observed in the actual reads of the JUN ChIP-seq data so it strongly suggests that our prediction that this candidate mutation creates a *de novo* AP1 binding site (**Figure 5**).

**Figure 5.**
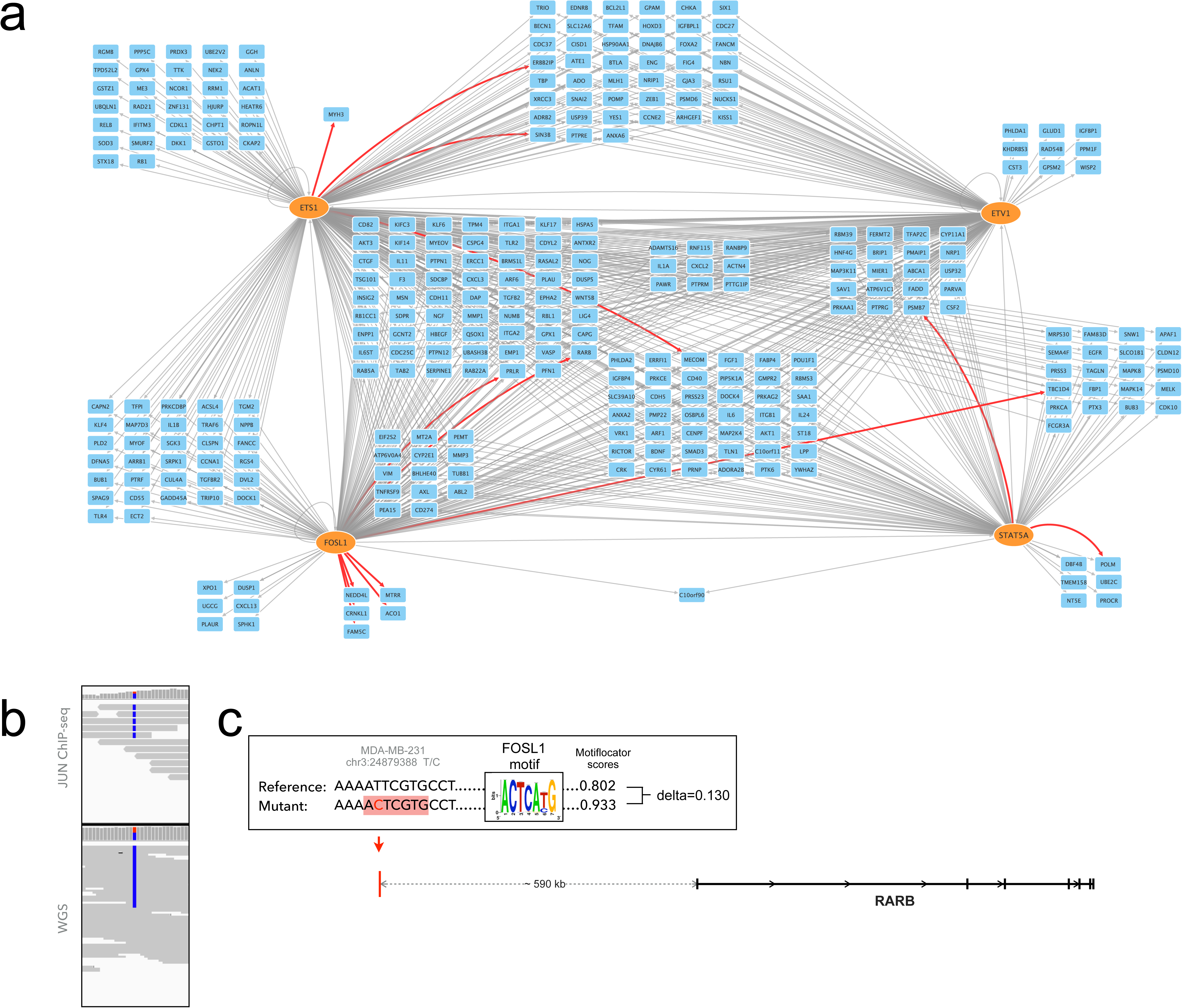
Personalized gene regulatory network of MDA-MB-231. Gene regulatory network inferred by motif-enrichment analysis from over-expressed genes (Z- score >=2) of breast cancer cell lines MDA-MB-231. Four master regulators are identified for this cell line: ETS1, ETV1, FOSL1, STAT5A. The grey edges represent the link between TFs and TGs based on iRegulon analysis, while red edges indicate gain of motifs caused by the *cis*-GoF. All the represented TGs are over-expressed in MDA-MB-231, are associated with breast cancer and are known cancer drivers. **b**) A *cis*-GoF mutation in a distal enhancer (590Kb upstream) of *RARB* gene creates a *de novo* AP1 binding site (resulting in a red edge between AP1 family transcription factor FOSL1 and *RARB*). **c**) IGV screenshot shows that the mutation is heterozygous in MDA-MB-231 whole genome sequence data (below), and homozygous in JUN ChIP-seq data (above).

Next, we investigated whether the predicted *cis*-regulatory mutations were present in an allele-specific manner in the expression data, in other words we checked if the mutation is associated to a gene with allele specific expression (ASE). Using coding heterozygous SNPs from WGS calls and RNA-seq data (which was not available for SKOV-3 and OVCAR-3) we identified genes with ASE, and this revealed that 154 of 468 candidate *cis-*GoF mutations show allelic bias in expression data (**Supplementary Table 6**). Note that our effort to identify mutations showing allelic bias in regulatory data failed since the coverage of H3K27Ac data was too low to determine variant allele frequency (80/98 candidate *cis*-GoF mutations in peaks have a depth of coverage below 5). When we expanded our search to also include SNPs, we identified 1304 SNPs in H3K27Ac peaks with a motif gain for a master TF, and of these, 41 show allelic bias in the regulatory data (**Supplementary Table 11**). This illustrates that gain of important motifs can yield allele-specific regulatory activity, but very few non-dbSNP, thus candidate somatic mutations were identified with this property across the 10 cell lines. On the other hand, by combining regulatory activity information and RNA-seq based ASE we found evidence of selection for 217 of 468 candidate *cis*-GoF mutations (**Figure 4**). In conclusion, *μ-cis*Target can be applied to matched genome-transcriptome data, or to matched genome-epigenome data, to obtain non-coding gain-of-function mutations resulting in gains of subtype-specific master regulators, near over-expressed oncogenes.

## Discussion

Whole-genome re-sequencing of cancer genomes is taking a prominent place in research and the clinic. The identification and prioritization of candidate driver mutations in the non-coding genome is therefore a key challenge. Indeed, several recent studies indicate an important role for *cis*-regulatory mutations in disease, not only in cancer (e.g., *TERT*, *TAL1* [3–5]), but also in complex diseases (e.g., Type II Diabetes FTO locus [43], Parkinson’s disease [44]) and in familial disorders (e.g. preaxial polydactyly [45]). However, other studies have highlighted that cancer *cis*-regulatory mutations are usually not recurrent across patients, except for a few exceptions such as the *TERT* promoter mutation. To reconcile these two opposing directions, we decided to step away from recurrence calculations in a cohort, but to work under the assumption that *cis*-regulatory mutations may be rarer than expected. A consequence of this assumption, if each sample harbours none or only few *cis*-regulatory mutations, is that statistical enrichment analyses may fail to provide meaningful results. Rather, the identification of functional *cis*- regulatory mutations may require a more *ad hoc* biological approach, which we explored in this study.

We are not the first to score and prioritize candidate mutations based on their putative gain (or loss) of transcription factor binding sites. In fact, most previously existing methods, such as FunSeq2, OncoCis, or RegulomeDB, use an *ad hoc* approach to annotate and filter candidate mutations based on motif loss/gain (FunSeq2, OncoCis) or on regulatory data from publicly available databases such as ENCODE (RegulomeDB). However, several important pieces of information are not utilized by previously existing tools, and are explored in our study. Firstly, the gain (or loss) of a motif is expected to be functionally relevant if the transcription factor itself is expressed in the cancer cells under study. If the transcription factor is (or was) not expressed, the gain of a binding site is not expected to be under positive selection.

Secondly, the motif gain (for an expressed transcription factor) should preferably represent a new binding site for a master regulator, meaning that this (over-expressed) transcription factor regulates other genes in the cancer cell *through this motif*. The approach we present here is to our knowledge the first one to take this criterion into account. Practically, to address this challenge we use a patient-specific, or subtype specific gene signature, in which over-represented motifs represent candidate motifs of master regulators.

Thirdly, the motif gain should yield either *de novo* regulatory activity of the encompassing enhancer, or should strengthen/amplify that enhancer. Such a gain of function may be visible as allele-specific bias of regulatory activity, whereby for example the ChIP-seq reads are homozygous for the variant.

Fourthly, the motif and enhancer gain should result in the up-regulation of a nearby oncogene, so that it provides a growth advantage to the cancer cell, and can be positively selected. Moreover, under this fourth piece of information, we expect that the predicted target gene is actually a known oncogene. Indeed, it is rather unlikely that previously unknown oncogenes could be discovered that are only up-regulated by a *cis*-regulatory mutation, and not by any other means (e.g., duplication, translocation, or mutation).

Our approach currently focuses on gain-of-function mutations that generate new binding sites for over-expressed activators, yielding up-regulation of a nearby oncogene. Clearly, several other scenarios are not covered by our proof-of-concept analyses. These include for example: the loss of an activator binding site near a tumour suppressor (e.g., loss of p53 binding site); the loss of a repressor binding site near an oncogene; or the gain of a repressor binding site near a tumour suppressor. We have focused in this study on the gain of an activator site near an oncogene because the currently known *cis*-regulatory driver mutations are all of this class, and this is the most conceivable and most pragmatic way to work within the context of motif discovery and gene regulatory networks. Future work is needed to address the other categories, and may reveal new types of *cis*-regulatory mutations. Another possible increase of sensitivity would be to also allow motif gains for *relevant* transcription factors that are expressed in the sample, and related to the cancer type, but for which the motif was not enriched in the input gene signature.

Our method provides a handle on understanding non-coding mutations in the context of regulatory genomics, thus we envisioned *μ-cis*Target not as a method for a final analysis but rather a starting point for in depth analysis. The typical use of a strategy like we depict here can be either to annotate a cancer genome with functional information regarding *cis*-regulatory mutations; or in a research context to generate a list of candidate mutations that can be further tested in a targeted screen, for example using massively parallel enhancer-reporter assays or CRISPR-Cas9 based modulation/mutation of the candidate mutations.

In conclusion, we present a computational framework inspired by a cancer biological viewpoint on oncogenic driver mutations. Our method can be used to identify candidate *cis*-regulatory mutations using sequence information alone, but works best on samples with combined genome and transcriptome data; while optimal results can be obtained if also matching epigenome data is available. Overall our results suggest the presence of only few *cis*-regulatory driver mutations per genome in cancer genomes that may alter the expression levels of specific oncogenes.

## Materials and methods

### Analysis of melanoma whole genomes for *TERT* promoter mutations

Expression data (Z-scores across all TCGA sequenced cancers) for seven melanoma samples (**Supplementary Table 1**) with *TERT* promoter mutations based on [22] were downloaded from Cosmic (v74). The signatures per sample (i.e. genes that have an expression Z-score above 1) were analysed by iRegulon to build the personalized gene regulatory networks (using the following parameters: motif collection 19K, putative regulatory region centered around TSS [20kb, 10kb, 500bp], motif rankings database across 10 and 7 species, NES threshold=3, ROC=0.03, rank threshold=5000).

Raw sequence data (bam files) for six melanoma samples was downloaded from GDC Data Portal (Legacy Archieve) and re-analyzed with VarScan [46] (command *somatic* with minimum variant allele frequency of 0.1, and a minimum coverage of 5 and 2 reads in tumor and normal samples, respectively). Non-coding mutations associated to melanoma-related or cancer driver genes were scored with MotifLocator for master TFs obtained in the previous step. The list of candidate *cis*- GoF mutations was filtered further using topologically associating domains (TADs) from 21 human cell lines and tissues [47].

### Whole-genome-sequencing on 10 cell lines

A549, COLO-205 and PC-3 cell lines were sequenced with Complete Genomics (CG), DU-145, OVCAR-3, SKOV-3 were sequenced with Illumina and HCT-116, HT-29, MDA-MB-231, and SK-MEL-5 were sequenced with both technologies. Complete Genomics sequencing was performed by the service provider using a proprietary sequencing-by-ligation technology. CG also performed primary data analysis, including image analysis, base calling, alignment and variant calling. CG variants were further filtered with depth of coverage threshold of 10, mutation coverage threshold of 5, and variant allele frequency threshold of 0.20. Illumina sequencing was done in accordance with the manufacturer's protocol. Primary data analysis was performed with manufacturer's software Casava (v1.8.2). Illumina variants were further filtered with depth of coverage threshold of 10 and mutation coverage threshold of 5. Variant calls were intersected for samples that were sequenced with both technologies. Coding variants were subtracted using protein coding exon locations from GENCODE (v19). Variants were annotated as non-SNPs or SNPs using dbSNP build 144 [48].

### Cell line specific regulatory data

H3K27ac ChIP-seq data for 6/10 cell lines were obtained from GEO and literature: A549 and HCT-116 (GSE31755) [49], HT-29 (GSE53602), PC-3 [50], MDA-MB-231 [51], SK-MEL-5 (GSE60666) [52]. For the remaining 4 cell lines (OVCAR-3, SKOV-3, DU-145, Colo-205), ChIP-seq was performed in this study. The cell lines were grown to ∼85% confluence per 15-cm dish. A total of 20 million cells per sample were collected, yielding ∼20 fractions of chromatin. ChIP samples were prepared following the Magna ChIP-Seq preparation kit using at least two chromatin fractions and 2–2.5?μg of antibody per fraction. Anti-histone H3 acetyl K27 antibody (ab4729, Abcam) antibody was used for ChIP. Per sample, 5–30?ng of precipitated DNA or input was used to perform library preparation according to the Illumina TruSeq DNA Sample preparation guide. In brief, the immunoprecipitated DNA was end-repaired, A-tailed and ligated to diluted sequencing adapters (1/100). After PCR amplification (15–18 cycles) and bead purification (Agencourt AmpureXp, Analis), the libraries with fragment size of 300–500?bp were sequenced using the HiSeq 2000 (Illumina). Sequence reads were mapped to the reference genome (hg19-Gencode v18) using Bowtie2 2.1.0 and narrow peaks were called using MACS2 algorithm (q-value < 0.001) [53]. Then the peaks less than 350 bp from each other were merged. Additionally, we have used peak calls from ChIP-Atlas database for 433 regulatory datasets (TF and chromatin ChIP-seq) across 7 cell lines (**Supplementary Table 10**, http://chip-atlas.org).

### Selection of cancer type specific TFs and TGs

Lists of cancer-type specific genes were extracted from NCBI Gene database (http://www.ncbi.nlm.nih.gov/gene) (**Supplementary Table 2**). Next, a collection of 1050 known cancer driver genes compiled from different resources [54–59] was used to further annotate the cancer type related genes (**Supplementary Table 2**).

### Cell line specific gene signatures and identification of sample-specific master regulators

Gene expression data (Z-scores) per cell line was obtained from Cosmic Cell Lines project (http://cancer.sanger.ac.uk/cell_lines). Cell line specific gene sets were created by selecting genes that have an expression Z-score above 1. The signatures were analysed by iRegulon to build the personalized gene regulatory networks (using the following parameters: motif collection 19K, putative regulatory region centered around TSS [20kb, 10kb, 500bp], motif rankings database across 10 and 7 species, NES threshold=3, ROC=0.03, rank threshold=5000). TFs that are cancer type related and expressed in the cell line (Z-score above 1) were considered as master regulators of the corresponding cell line (**Table 1**).

### Detection of candidate mutations

The non-coding mutations were assigned to genes using GREAT tool (up to 1 Mb) [60]. Only the non-coding mutations associated with genes that are

a. over-expressed in the cell line (expression Z-score above 1),
b. relevant to the cancer type <u>or</u> on the list of known cancer drivers were scored by MotifLocator [13].

All the motifs corresponding to master TFs from the personalized gene regulatory networks were tested to reveal if any mutation from the matched cell line cause a gain of any of these motifs.

### Annotation of mutations using topologically associating domains (TADs)

The assignment of the mutations to the potential target genes within the space 1 Mb is further annotated using a large dataset of TAD boundaries generated for 21 samples (14 human tissues and 7 human cell lines) [47]. In the output file, the information whether the mutation and the gene fall between the boundaries of the certain sample is provided, i.e. if the mutation is in the same TAD region with the promoter of the gene. This annotation can be used to further filter mutation-to-genes associations.

### Genome-wide screening of DNA sequences by MotifLocator

A collection of 8053 unique PWMs directly annotated to 1628 TFs was used for scoring with MotifLocator [13]. Scoring was performed at the mutation sites with a window size between 20 bp to 60 bp depending on the motif size (20 bp if the motif size is 15 bp or less, 30 bp if motif size is between 16 bp and 25 bp, and 60 bp if the motif size is larger than 25 bp). The variants with mutant MotifLocator score >= 0.90 and delta >= 0.1 were selected (where delta represents the difference between MotifLocator scores of mutant and wild type sequences).

### Validation of candidate mutations using matched transcriptome and epigenome data

RNA-seq data (bam files) was downloaded for 8 (A549, COLO-205, DU-145, HCT-116, HT-29, MDA-MB-231, PC-3, SK-MEL-5) cell lines from GDC Data Portal (Legacy Archieve). Heterozygous SNPs per cell line were obtained from whole genome mutation calls by requiring at least 5 reads for reference and variant allele, as well as requiring at least 0.10 VAF. Heterozygous SNPs were intersected for samples that were sequenced with both Illumina and Complete Genomics. Allele specific expression per cell line was calculated using MBASED one-sample analysis [61]. Any gene with an adjusted p-value ≤ 0.05 and estimated MAF ≥ 0.6 was annotated as exhibiting allele-specific expression.

## Availability

The data generated for this study (H3K27ac ChIP-seq for OVCAR-3, SKOV-3, DU-145, COLO-205) will be deposited in NCBI’s Gene Expression Omnibus. Genome data are being deposited at the European Genome-phenome Archive (EGA, http://www.ebi.ac.uk/ega/) which is hosted at the EBI. The code for *μ-cis*Target is freely available at https://github.com/aertslab/mucistarget and a web-implementation is available at http://mucistarget.aertslab.org.

## Acknowledgements

This work is funded by The Research Foundation - Flanders (FWO, www.fwo.be) (G.0791.14), Special Research Fund (BOF) KU Leuven (http://www.kuleuven.be/research/funding/bof/) (grant PF/10/016), Foundation Against Cancer (http://www.cancer.be) (2012-F2 to SA). ZKA is funded by a research grant from Kom of tegen kanker. HI has a PhD Fellowships from the agency for Innovation by Science and Technology (IWT, www.iwt.be). The funders had no role in study design, data collection and analysis, decision to publish, or preparation of the manuscript.

## Competing interests

The authors declare that they have no competing interests.

## Supplementary Tables

**Supplementary Table 1.** Analysing 7 melanoma whole genomes with *TERT* promoter mutation using *μ-cis*Target

**Supplementary Table 2.** Cell type specific genes and potential driver genes

**Supplementary Table 3.** Candidate *cis*-GoF mutations in 3 melanoma whole genomes

**Supplementary Table 4.** Transcription factor ChIP-seq track enrichment for predicted master regulators

**Supplementary Table 5.** Overview of variants in 10 cell lines; sequencing technologies, number of variants before and after various filtering steps.

**Supplementary Table 6.** Candidate (non-SNP) *cis*-GoF mutations in 10 cell lines.

**Supplementary Table 7**. Protein altering coding mutations in genes affected by a *cis-*GoF mutation

**Supplementary Table 8**. Genes affected by *cis*-GoF mutations across 10 cell lines

**Supplementary Table 9.** Enrichment of *cis*-GoF mutations and SNPs in active regulatory regions

**Supplementary Table 10**. Matching TF and chromatin ChIP-seq data for 10 cell lines from ChIP-Atlas database

**Supplementary Table 11.** *cis*-GoF SNPs with regulatory allelic bias in 10 cell lines

## Supplementary Figures

**Supplementary Figure 1.** i-*cis*Target motif enrichment results for the MYB ChIP-seq peaks on JURKAT. Distribution of the area under the curve results for all 18832 motifs tested using the top 500 MYB ChIP-seq peaks on JURKAT cell line as input set. The arrows indicate the first motif found for RUNX and MYB transcription factors together with their respective normalized enrichment scores (NES). The table shows the top ten enriched motifs with their respective origins, NES and motif logos.

**Supplementary Figure 2.** Heatmap visualizing pathways each master regulator per cell line involved based on GeneAnalytics analysis (geneanalytics.genecards.org) [63]

**Supplementary Figure 3.** Scatter plot showing the number of *somatic* coding mutations in 10 cell lines versus number of *cis*-GoF mutations. Here the *somatic* coding mutations were extracted from Cosmic Cancer Cell Lines project database by selecting coding mutations with “*Confirmed somatic variant”* or “*Reported in another cancer sample as somatic”* tags. The black line represents the linear fit excluding the colon cancer cell line HCT-116 and dashed grey line represents the linear fit including all cell lines.

